# Autocrine Proteinase Activated Receptor (PAR) mediated signaling in prostate cancer cells

**DOI:** 10.1101/2022.08.22.504840

**Authors:** Arundhasa Chandrabalan, Rithwik Ramachandran

## Abstract

Proteinase activated receptors (PARs) are G protein-coupled receptors (GPCRs) activated by limited N-terminal proteolysis. A variety of proteolytic enzymes derived from the coagulation cascade and inflammatory milieu activate PARs, however specific activators in different physiological and pathophysiological contexts remain poorly defined. PARs are highly expressed in many cancer cells and regulate various aspects of tumor growth and metastasis. Endogenous proteinases that regulate PARs in the setting of various tumors however remains unresolved. Prostate cancer (PCa) remains a major cause of mortality in men despite advances in early detection and clinical intervention. PAR expression has been reported in PCa, however, their role here remains poorly defined. In androgen independent PC3 cells, we find functional expression of PAR1 and PAR2 but not PAR4. Using genetically encoded PAR cleavage biosensors, we find that PCa cells secrete proteolytic enzymes that cleave PARs and trigger autocrine signaling. Deletion of PAR1 and PAR2 using CRISPR/Cas9 combined with microarray analysis revealed genes that are differentially regulated by this autocrine signalling mechanism. Interestingly, several genes that are known PCa prognostic factors or biomarker were differentially expressed in PAR1-KO and PAR2-KO PC3 cells. We also examined PAR1 and PAR2 regulation of PCa cell proliferation and migration using PAR1 and PAR2-KO PC3 cells, as well as PAR1 and PAR2 specific agonists and antagonists. We find that PAR1 and PAR2 have opposite effects on PC3 cell proliferation and migration. In summary, we have identified an autocrine signaling mechanism through PARs as a regulator of PCa cell function.

## INTRODUCTION

Prostate cancer (PCa) is a major cause of mortality in men despite advances in early detection and clinical intervention. The heterogeneity of PCa makes it a challenge in identifying effective biomarkers and treatment. The exact mechanism driving the disease remains elusive, where understanding of PCa associated molecular signalling pathways would help us better understand the underlying mechanisms of the disease progression and would potentially aid in early diagnosis and/or prognosis.

The tumor microenvironment has emerged as a crucial factor in regulating cancer progression besides genetic background of the cells. Proteinases are vital component in the tumor microenvironment that regulates several functionally distinct processes in almost every hallmark of tumorigenesis^1^. Cancer progression is often linked with dysregulation of normal proteolysis regulating mechanisms, which leads to numerous proteinases with dramatically altered mostly upregulated activity and proteolytic remodelling of the microenvironment^2, 3^. Mainly proteinases including matrix metalloproteinases (MMP-1, -14)^2, 4^, tissue kallikreins (KLK-3, -6)^5^, cathepsins (CTS-B, -L, -S), matriptase, hespin, and urokinase plasminogen activator (uPA) mediate extracellular matrix (ECM) degradation resulting in cancer invasion and metastasis^2,6–9^. Of these proteinases, MMPs are well-known to have a role in ECM remodelling and several crucial roles in normal and pathological processes including inflammation, atherothrombotic disease and angiogenesis. MMPs have been extensively studied in cancer biology and found to possess direct signalling capabilities. However, in clinical trials, use of broad-spectrum MMP inhibitors to treat different types of cancer failed to be feasible, highlighting important roles MMPs play in normal physiology and the many redundant enzymes that can be present in pathology^10, 11^. In addition to promoting tumorigenesis, proteinases play a role in tumor suppression as well as in signalling pathways^12^. Evidently, proteinases associated tumor pathogenesis are highly complex^13, 14^. An approach that can modulate specific proteinase possibly in combination with secondary target may therefore be therapeutically effective.

Proteinase-activated receptors (PARs, PAR1-4) are a unique four-member family of G protein coupled receptors (GPCRs) activated by proteinases by limited receptor proteolysis and unmasking of a cryptic tethered ligand^15, 16^. Overexpression of PARs have been observed in various cancer cell types and have been implicated in many aspects of tumorigenesis associated cellular processes such as cell migration, invasion, metastasis, proliferation and angiogenesis^17–19^. Of the PARs, the function of PAR1 and PAR2 have been studied in a number of human cancer cells and or tissues,^20–25^ while the role of PAR3 and PAR4 is mostly unknown. PAR1 and PAR2 are mainly activated by serine proteinases, including thrombin, coagulation factors (FXa, FVIIa), activated protein C, trypsin, neutrophil enzymes and kallikrein-related peptidases, metalloproteinases (MMPs) and cysteine proteinases such as cathepsins and calpain, which are involved in cell signalling, cell chemotaxis, inflammation, vascular integrity and ECM remodelling^16, 26^. Remarkably, these proteinases are present in the tumor microenvironment^11^, and PAR1 and PAR2 signals to several of these proteinases promoting crosstalk between tumor cells and the microenvironment^24,27,28^. In addition, PAR1 has been recognized as an oncogenic protein under pathophysiological conditions in carcinoma that generally promotes cancer cell migration, invasion, metastasis and survival^25^. PAR2 and its activating proteinases are also overexpressed in various cancers and shown to be involved in angiogenesis (hypoxia-induced), cell proliferation and metastasis^29^. Though it is evident that PAR1 and PAR2 are implicated in pathophysiological processes in cancer progression and these receptors have emerged as promising therapeutic targets associated to metastasis, the precise mechanism of PARs downstream signalling that differentially regulate tumorigenesis has not been fully elucidated yet and warrants further investigation.

The identification of the ability of multiple proteinases to activate the PARs has expanded considerably the context within which these receptors might regulate physiological and pathophysiological responses in the body^15,30,31^. There has also been work from our group and others that have established the ability of different enzymes to trigger distinct intracellular signaling pathways downstream of PAR, which is in keeping with the reported ability of a number of GPCRs to exhibit such agonist biased signalling^16^. We recently reported that PARs exhibit biased signaling with the enzyme processing the receptor dictating the second messenger coupling^32, 33^. This interesting finding has led us and others to identify a number of proteinases that irreversibly cleave PARs, but do not couple to the same second messenger pathways^34^. This in turn makes it hard to efficiently identify endogenous receptor cleaving enzymes. In order to enable a rapid investigation of an enzyme ability to cleave the receptor, without relying on second messenger coupling assays, in this study we used genetically encoded reporter constructs that enable direct monitoring of receptor cleavage. In addition, here we used PAR knockout PCa cell lines and microarray analysis to explore differentially expressed oncogenes associated in PCa.

## MATERIALS AND METHODS

### Materials

Thrombin (human plasma, high activity) and trypsin (porcine pancreas, Type IX-S, 13000-20000 BAEE units/mg protein) was purchased from Millipore Sigma. PARs activating peptides TFLLR-NH_2_ (PAR1), SLIGRL-NH_2_ (PAR2) and AYPGKF-NH_2_ (PAR4) (≥95% purity by HPLC) were synthesized by GenScript. PAR1 selective antagonist vorapaxar was purchased from Adooq Bioscience, and PAR2 antagonist AZ3451 was from Sigma. The stock solutions of the PAR antagonists were prepared in dimethyl sulfoxide (DMSO, BioShop). The broad-spectrum serine proteinase inhibitor 4-(2-aminoethyl)benzenesulfonyl fluoride hydrochloride (AEBSF) was obtained from ThermoFisher Scientific, and soybean trypsin inhibitor (STI) was from Millipore Sigma. The broad-spectrum MMP inhibitor batimastat (BB-94, ≥98%) was purchased from Millipore Sigma. The thrombin selective inhibitor D-phenylalanyl-L-prolyl-L-arginine chloromethyl ketone dihydrochloride (PPACK.2HCl) was from Millipore Sigma. The stock solutions of the agonists were made in 25 mM 4-(2-hydroxyethyl)-1-piperazine ethanesulfonic acid (HEPES, Fisher Scientific) and the enzyme inhibitors were reconstituted according to manufacturers’ instructions. All samples were diluted to appropriate working concentrations in Hanks’ Balanced Salt Solution containing CaCl_2_ and MgCl_2_ (HBSS, Gibco, ThermoFisher Scientific).

### Cell culture

Human Prostatic Cancer cells PC3 (ATCC_®_CRL-1435) were cultured in Ham’s F-12K (Kaighn’s) Medium supplemented with 1 mM L-Glutamine, 100 U ml^-1^ penicillin, 100 µg ml^-1^ streptomycin, 1 mM sodium pyruvate, and 10% v/v heat inactivated Fetal Bovine Serum (FBS) from Gibco^®^ ThermoFisher Scientific. Chinese Hamster Ovary cells (CHO-K1, Sigma) stably expressing either pcDNA3.1(+) nLuc-PAR1-eYFP or nLuc-PAR2-eYFP (nLuc, nano-luciferase; eYFP, enhanced yellow fluorescent protein) constructs were cultured in Ham’s F-12 (1×) Nutrient Mix (Gibco ThermoFisher Scientific) with 1 mM L-Glutamine, 100 U ml^-1^ penicillin, 100 µg ml^-1^ streptomycin, 1 mM sodium pyruvate, 10% v/v FBS, and 600 µg ml^-1^ geneticin selective antibiotic (G418 Sulfate, Gibco_®_ ThermoFisher Scientific). The cells were grown in a T75 cell culture flask (Nunc, Fisher Scientific) in a humidified cell culture incubator with 5% CO_2_ at 37 °C. Cells at ∼80-90% confluency were detached with phosphate buffered saline (PBS, Gibco® ThermoFisher Scientific) solution supplemented with 1 mM EDTA (Fisher Scientific), centrifuged at 180 × *g* for 5 minutes, and sub-cultured as needed.

### CRISPR/Cas9 PAR knockout (KO) PC3 cells

PC3 cells endogenously express PAR1 and PAR2. PAR1 deficient (PAR1-KO) and PAR2 deficient (PAR2-KO) PC3 cell lines were generated using CRISPR/Cas9 targeting with targeting strategy and procedures essential as described previously^35, 36^. Briefly a specific guide targeting PAR1 (GATAGACACATAACAGACCG) or PAR2 (CCCCAGCAGCCACGCCGCGC) was cloned into the lentiCRISPR v2 plasmid (Addgene plasmid # 52961; a kind gift from Dr. Feng Zhang) and transfected into PC3 cells. 48 hours after transfection cells were placed in media containing puromycin (5 μg/ml) and knockout cell lines were established by dilution cloning and screening of functional response to PAR1 or PAR2 specific agonists using a calcium signaling assay. Final cell lines used in all experiments were verified to be deficient in PAR1 or PAR2 while retaining activity to the non-targeted PAR receptor.

### Calcium signalling

PAR activated calcium release was measured as described previously with some modifications^37^. WT, PAR1-KO and PAR2-KO PC3 cells were plated in a 96-well black cell culture microplate (Optical Bottom, Polystyrene, Thermo Scientific) at a density of 1 × 10^4^ cells per well and cultured for 48 h in Ham’s F-12K complete culture medium. The cells were rinsed with PBS (2 × 100 µl per well) and incubated with 50 µl per well fluorescent calcium indicator Fluo-4 NW (λ_Ex/Em_ 494/516 nm, catalogue no. F36206, Life Technologies) at 37 °C for 30 minutes and for further 15 minutes at room temperature in the dark. Ca^2+^ influx was indicated by a change in fluorescence measured on a FlexStation3 (Molecular Devices) microplate reader. The real-time fluorescence spectra were recorded using a Flex Fluorescence Bottom-read protocol with an excitation and emission wavelengths of 494 and 515 nm, respectively. The total run time of each spectrum was 180 s, with baseline recorded for 20 s prior to addition of agonists. To measure antagonism, the cells were pre-treated with antagonists for 30 minutes at RT before measuring agonist responses as above. Calcium ionophore (A23187, 6 µM, Sigma-Aldrich) was used as a control in all experiments to determine the maximum calcium response possible in each plate of the cells.

### PC3 cells supernatant

PC3 cells were plated in a 100 mm × 20 mm cell culture dish (Falcon, Corning) at a density of 10 × 10^4^ cells/ml and cultured in F-12K complete medium for 24 h. The cells were rinsed with PBS (2 × 4 ml), and 4 ml of serum-free and phenol-red free Eagle’s Minimum Essential Medium (EMEM, Mod. 1×) was added and cultured for 24 or 48 h. PC3 supernatants (conditioned media) were collected, centrifuged at 180 × *g* for 5 minutes to remove cell debris and frozen at -80 °C until use in the assays.

### Luciferase assay

CHO-K1 cells stably transfected with nLuc-PAR1/2-eYFP constructs were plated in a 96-well cell culture plate (polystyrene, flat-clear bottom, Nunclon Delta, Nunc, ThermoFisher Scientific) at a cell density of 1 × 10^4^ cells per well and cultured for 48 h in F-12 complete culture medium. The cells were rinsed with HBSS (3 × 100 µl) and incubated with 100 µl HBSS at 37 °C for 15 minutes. 50 µl of cell supernatant was transferred to a 96-well white microplate (polystyrene, Nunclon Delta, Nunc, ThermoFisher Scientific) to measure the basal luminescence. The cells were incubated with 50 µl test samples or controls at 37 °C for 15 minutes, and 50 µl of cell supernatant from each well was aliquoted as before. The test samples (PC3 supernatant) were untreated or pretreated with a proteinase inhibitor BB-94 (25 µM), AEBSF (100 µM), PPACK (1 µM) or STI (100 µg ml^-1^), separately, by incubating at room temperature for 30 minutes. Nano-Glo Luciferase Assay Substrate furimazine (2 µl ml^-1^, Promega) was added and the nLuc released upon proteolytic cleavage of the receptor was measured on a plate reader Mithras LB 940 (Berthold Technologies, measurement time: 1 s per well).

### Cell migration

PC3 cell migratory properties were measured using a wound healing scratch assay approach. WT, PAR1-KO and PAR2-KO PC3 cells were plated in a 12-well cell culture plate (Greiner Bio-One) at a density of 40 × 10^4^ cells/ml per well and cultured for 24 h in F-12K complete culture medium to form a monolayer. The cells were rinsed with PBS (2 × 0.5 ml per well) and the scratch was made in the middle of the well from top to bottom using a P200 pipette tip having the cells in 0.5 ml PBS. The cells were rinsed with PBS (3 × 0.5 ml per well), and 1 ml low-serum (0.1% FBS) F-12K culture medium was added. Low serum culture medium was used to suppress cell proliferation. The cells were untreated or treated with a PAR agonist or antagonist(s) at a concentration that elicits a maximum response in calcium signalling. Widefield images were taken immediately (0 h) and every 6-12 h for 48 h at magnification 10× using a Nikon Eclipse Ti2 inverted microscope with NIS Elements AR software. Cell migration rate or scratch area closure was analyzed using an open-source image processing program Fiji-ImageJ, in which the cell-free area over time was measured. The rate of cell migration was represented as a percentage of change in the scratch area over time.

### Cell proliferation

PC3 cell proliferation was quantified by an MTT based colorimetric assay using the Cell Proliferation Kit I (Roche, Sigma Aldrich). WT, PAR1-KO and PAR2-KO PC3 cells (100 µl per well) were plated in a 96-well cell culture plate (polystyrene, clear flat-bottom, Nunclon Delta, Nunc, ThermoFisher Scientific) at a cell density of 2 × 10^3^ cells per well and cultured for 24 h in F-12K complete (10% FBS) culture medium. The culture medium was replaced with low-serum (2%) F-12K medium and incubated for 3 h in the cell culture incubator. The cells were untreated or treated with a PAR agonist or antagonist diluted in low-serum (2%) F-12K medium at a concentration that elicits a maximum response in calcium signalling, and incubated for 24, 48, 72 and 96 h. MTT labeling reagent (10 µl) was added and incubated for 4 h. Solubilization solution (100 µl) was added to the MTT treated wells and incubated overnight. The absorbance at 570 nm was measured on a plate reader (FlexStation3).

### Microarray

Total RNA from WT, PAR1-KO and PAR2-KO PC3 cells were extracted using a RNeasy Mini Kit (Qiagen). RNA quality, RNA Integrity Number (RIN), were measured using an Agilent RNA Nano Chips on a 2100 Bioanalyzer (Agilent Technologies). 250 ng of total RNAs with a RIN of ≥8 were labelled using the GeneChip Whole Transcript Express Labeling Assay and hybridized to GeneChip Mouse Gene 2.0 ST Arrays (Affymetrix) at 45 °C for 16 h (SACRI Microarray and Genomics Facility, University of Calgary). Arrays were stained and washed with GeneChip Fluidics Station 450 and scanned on a GeneChip Scanner 3000 7G (Affymetrix). Probe cell intensity data CEL files were generated on GeneChip Command Console (Affymetrix) software. The PrimeView array gene expression analyses were performed on Transcriptome Analysis Console (TAC 4.0, Affymetrix) software. Robust Multi-Array Average (RMA) summarization method was used to log 2 transform and normalize the data. Differentially expressed genes with <-2 and >2 fold-change were filtered and associated signalling pathways were identified by WikiPathways as well as manual extraction. Raw microarray data files were deposited in the NCBI Gene Expression Omnibus (GEO) repository with accession number GSE211813.

### Graphical and Statistical analyses

The release of intracellular calcium [Ca^2+^]^i^ measured as a change in relative fluorescence unit (ΔRFU) in calcium signalling assays were normalized by subtracting its baseline. The luminescence measurements in the luciferase assay were normalized by subtracting the basal luminescence signal obtained with HBSS. The calcium signalling and luciferase assay data represents the mean of at least three independent experiments (*N* ≥ 3), performed either in duplicates (*n* = 6) or triplicates (*n* = 9), with their standard error of the mean (SEM). The concentration-effect curves were plotted and analyzed using the standard slope (1.0) dose-response stimulation three parameters model with a non-linear regression curve fit in GraphPad Prism 8, and the values of logEC_50_ were obtained with their SEM. To determine statistical significance (*p* < 0.05) in potency and efficacy, the extra-sum-of-squares F-test in GraphPad Prism 8 was utilized. Statistical significance between non-treated and treated samples were determined with Bonferroni-Dunn multiple comparisons t-tests at *p* < 0.05.

## RESULTS

### Characterization of PC3 cells

Functional PAR1, PAR2 and PAR4 expression in PC3 cells was examined through monitoring Gα_q/11_-mediated calcium signalling in response to canonical PAR activating enzymes (PAR1 and PAR4: thrombin, PAR2: trypsin) and PAR-specific agonist peptides (PAR1: TFLLR-NH_2_, PAR2: SLIGRL-NH_2_, PAR4: AYPGKF-NH_2_) (**Figure 1** and **S1**). In WT-PC3 cells, a calcium signalling response was evident in cells treated with thrombin (0.03 – 10 U ml^-1^, EC_50_ 0.4 U ml^-1^, **Figure 1A**), trypsin (0.3 – 100 nM, EC_50_ 5.6 nM, **Figure 1C**), and the PAR1 and PAR2 activating peptides TFLLR-NH_2_ (1 – 300 µM, EC_50_ 6.6 µM, **Figure 1B**) and SLIGRL-NH_2_ (1 – 300 µM, EC_50_ 3.9 µM, **Figure 1D**), respectively. However, the PAR4-specific agonist peptide AYPGKF-NH_2_ (1 – 300 µM, **Figure S1-E**) did not trigger signalling. In PAR1-KO PC3 cells, thrombin and the PAR1-specific agonist TFLLR-NH_2_ did not activate signalling, except at the highest concentration tested (**Figure 1A**, **B**). Both thrombin and TFLLR-NH_2_ activated the endogenously expressed PAR1 in a concentration-dependent manner in PAR2-KO PC3 cells (thrombin EC_50_ 0.2 U ml^-1^, TFLLR-NH_2_ EC_50_ 2.3 µM, **Figure 1A**, **B**). This further confirmed that thrombin signals through PAR1 in WT-PC3 cells and not through PAR4. The thrombin and TFLLR-NH_2_ responses seen in PAR1-KO cells is due to PAR2 activation that can occur at high concentration for both of these activators.

**Figure 1:**
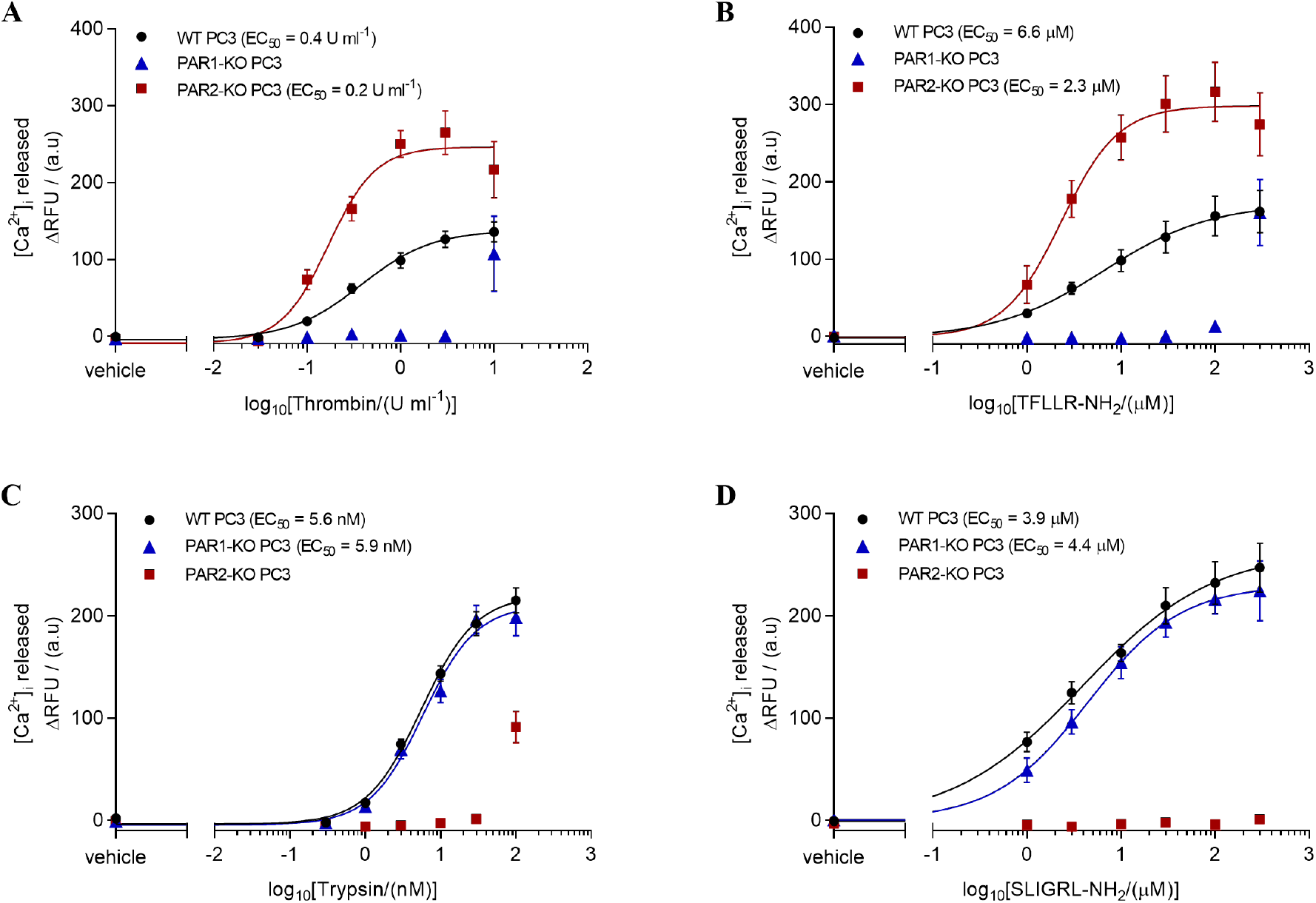
PAR1 and PAR2 are functionally expressed in WT-PC3 cells and are absent in KO cells. PC3 cells were characterized for PAR expression through Gαq/11-coupled calcium signalling with known PAR1 agonists (**A**) thrombin (0.03 – 10 U ml^-1^) and (**B**) TFLLR-NH2 (1 – 300 µM), and with PAR2 agonists (**C**) trypsin (0.3 – 100 nM) and (**D**) SLIGRL-NH2 (1 – 300 µM) on WT, PAR1-KO, and PAR2-KO PC3 cells. Each data point represents the mean ± SEM (N = 3). Extra-sum-of-squares F-test was utilized to compare significant (*p* < 0.05) differences in potency and efficacy between WT and PAR-KO PC3 responses.

Trypsin and the PAR2-specific agonist SLIGRL-NH_2_ did not show a response in PAR2-KO PC3 cells (**Figure 1C**, **D**), except with trypsin at the highest concentration tested due to activation of PAR1. Both PAR2 agonists triggered a concentration-dependent response in PAR1-KO PC3 cells (trypsin EC_50_ 5.9 nM, SLIGRL-NH_2_ EC_50_ 4.4 µM, **Figure 1C**, **D**). Overall, these data demonstrate that PAR1 and PAR2 are functionally expressed in PC3 cells while PAR4 is not (**Figure S1**). In addition, a significant increase in thrombin and TFLLR-NH_2_ responses are seen in PAR2-KO PC3 cells (**Figure 1A**, **B**) indicating that PAR1 expression is upregulated when PAR2 is knocked out in PC3 cells.

### PAR cleaving proteinases in PC3 cells

Genetically encoded PAR biosensor nLuc-PAR1-eYFP and nLuc-PAR2-eYFP, stably transfected in CHO cells, allowed us to measure PAR-cleaving proteinase activity in PC3 cell-derived supernatants by monitoring the cleavage of the nLuc-tag from PAR N-terminus following protease cleavage of the receptor (**Figure 2**). CHO cells were chosen as the background for the reporter constructs since they do not produce significant amounts of endogenous PAR cleaving enzymes and their firm attachment to culture plates prevented contamination of the luciferase signal in the supernatant by detached cells. As expected, the standard PAR1 and PAR2 activating enzymes, thrombin (0.003 – 10 U ml^-1^) and trypsin (0.3 – 100 nM) showed a concentration-dependent PAR cleavage in nLuc-PAR1-eYFP (**Figure 2A**) and nLuc-PAR2-eYFP (**Figure 2B**) CHO cells, respectively.

**Figure 2:**
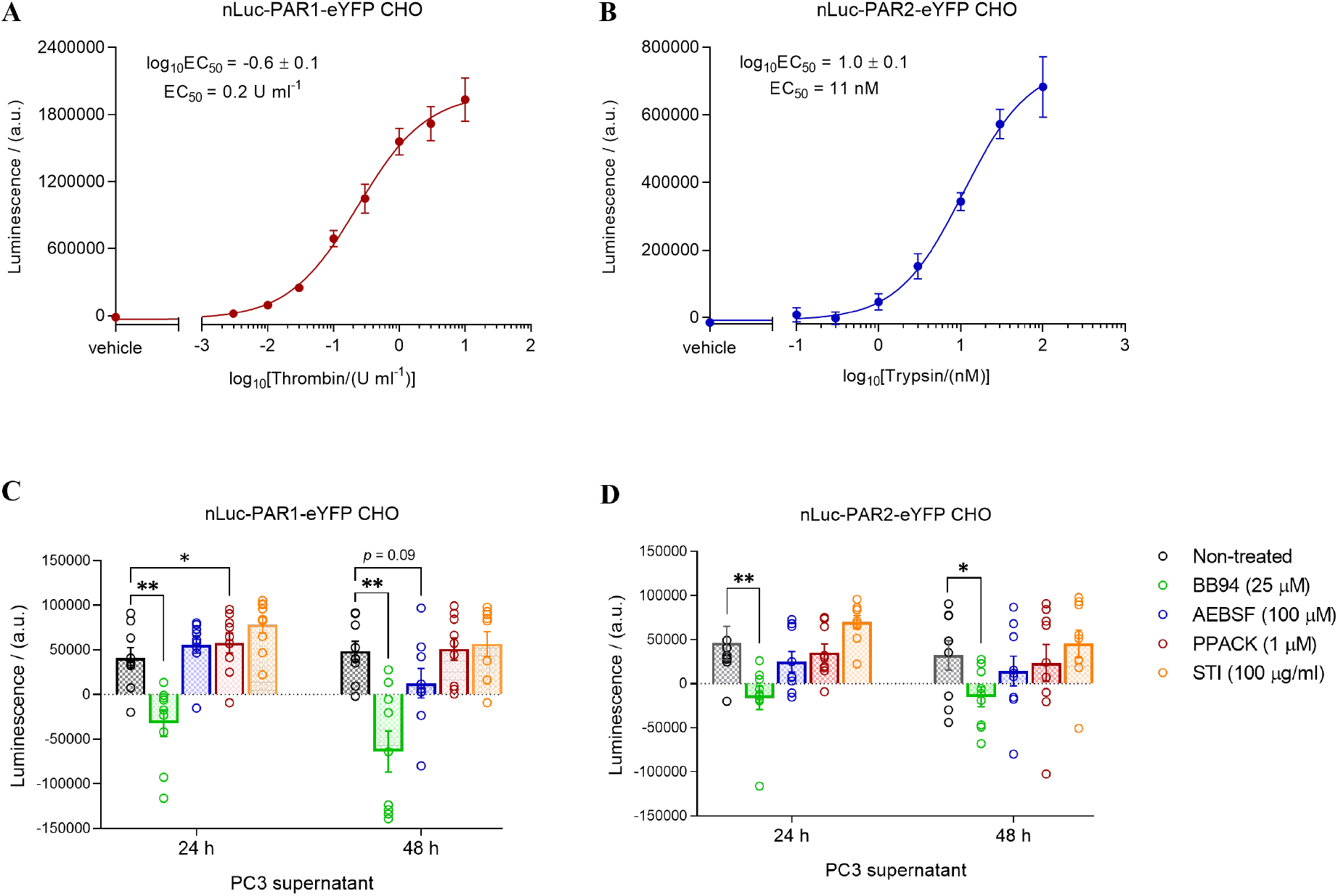
Identification of PAR1 and PAR2 cleaving enzymes in PC3 cells supernatant. Characterization of (**A**) nLuc-PAR1-eYFP and (**B**) nLuc-PAR2-eYFP transfected CHO cells with PAR1 and PAR2 agonists thrombin (0.003 – 10 U ml^-1^) and trypsin (0.1 – 100 nM), respectively. Cleavage of PARs expressed in CHO cells, (**C**) nLuc-PAR1-eYFP and (**D**) nLuc-PAR2-eYFP, by proteinases in PC3 cells cultured (24 and 48 h) supernatant was measured untreated or treated with a proteinase inhibitor BB94 (MMP), AEBSF (serine proteinase), PPACK (thrombin) or STI (trypsin). Each data point represents the mean ± SEM (N = 3). Bonferroni-Dunn multiple t-tests was utilized to determine statistical significance at \**p* < 0.05 and \*\**p* < 0.01 relative to the untreated control.

In order to examine whether PC3 cells produced PAR cleaving enzymes, conditioned media of cultured (24 or 48 h) PC3 cells were incubated with each of the biosensor cell lines. Surprisingly, cleavage of both PAR1 (**Figure 2C**) and PAR2 biosensors (**Figure 2D**) was evident in cells incubated with PC3 conditioned media. In order to determine the protease type present in PC3 supernatants we compared PAR1 and PAR2 cleavage following incubation with MMP and serine protease inhibitors. Interestingly the supernatants showed differential inhibition of PAR cleaving ability with broad spectrum MMP inhibitor (BB-94) and serine proteinase inhibitors (AEBSF and PPACK) (**Figure 2C**, **D**). A complete inhibition of both PAR1 and PAR2 cleavage was observed with MMP inhibitor BB-94 pretreated PC3 supernatants derived from 24 and 48 h culture (**Figure 2C**, **D**). A partial inhibition of PAR1 and PAR2 cleavage was seen with serine proteinase inhibitors AEBSF and PPACK pretreated supernatants (**Figure 2C**, **D**), PPACK showing a greater effect in inhibiting cleavage of PAR1 (**Figure 2C**). STI (trypsin-selective inhibitor) pretreated of supernatants did not inhibit PAR1 or PAR2 cleavage (**Figure 2C**, **D**). We also noted a decrease in PAR1 and PAR2 cleavage when comparing 24 and 48 h culture derived supernatants. This could be due to enzyme degradation over time in culture or inhibition of enzymes by endogenous proteinase inhibitors.

### PARs modulate PC3 cell migration

To determine the role of endogenously expressed PARs in PCa, the cell migratory property of PC3 cells was assessed in serum-starved WT, PAR1-KO and PAR2-KO PC3 cells by a scratch assay over 48 h period (**Figures 3** **and S2**). Remarkably, we found that the rate of cell migration is significantly enhanced in PAR1-KO PC3, while no significant differences were seen in PAR2-KO PC3, relative to the WT PC3 cell migration rate (**Figure 3A**). This suggests that in PC3 cells signalling downstream of PAR1 activation by cell secreted enzymes acts as a negative regulator of cell migration.

**Figure 3:**
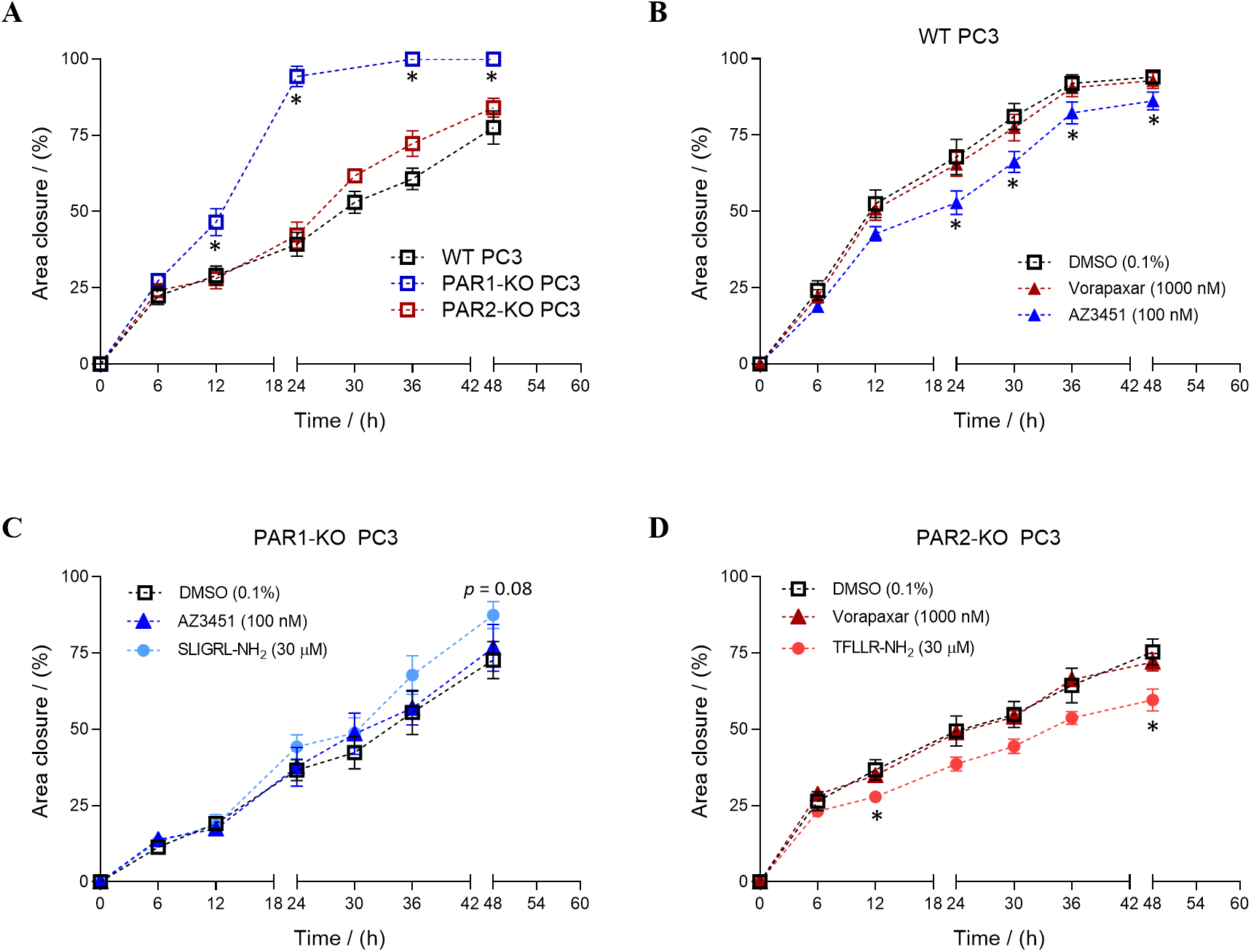
PARs regulate PC3 cell migration. PC3 cell migratory property assessed through a scratch assay over 48 h period in (**A**) untreated WT, PAR1-KO and PAR2-KO PC3, (**B**) PAR1 and PAR2 antagonists vorapaxar and AZ3451, respectively, treated WT-PC3, (**C**) PAR2 antagonist and agonist SLIGRL-NH2 treated PAR1-KO PC3, and (**D**) PAR1 antagonist and agonist TFLLR-NH2 treated PAR2-KO PC3 cells. Each data point represents the mean ± SEM (N ≥ 3). Bonferroni-Dunn multiple t-tests was utilized to determine statistical significance at \**p* < 0.05 relative to the (untreated) control.

In order to further examine the regulation of PCa cell migration by PARs, WT, PAR1-KO and PAR2-KO PC3 cells were treated with either PAR1 or PAR2 specific agonist or antagonist and the cell migration was similarly examined over 48 h (**Figure 3B**-**D**). Concentration of PAR1 and PAR2 antagonists for use in these assays was established through their ability to block thrombin and trypsin mediated calcium signalling on PC3 cells respectively (**Figure S3**). Interestingly, inhibition of PAR2 in WT PC3 cells with AZ3451 significantly inhibited cell migration while the PAR1 antagonist vorapaxar was without effect (**Figure 3B**). In PAR1-KO PC3 cells, PAR2 activation or inhibition did not affect cell migration (**Figure 3C**) while in PAR2-KO PC3 cells activation of PAR1 with TFLLR-NH_2_ in cells significantly lowered the migration rate (**Figure 3D**), but no change was observed with the PAR1 antagonist vorapaxar. Together these data suggest that PAR1 inhibits and PAR2 promotes cell migration in PCa, though unexpectedly the receptor KO cells, and antagonist treatment did not produce identical responses.

### PARs modulate PC3 cell proliferation

The role of endogenously expressed PARs in regulating PCa cell proliferation was assessed using an MTT assay in WT, PAR1-KO and PAR2-KO PC3 cells over 24 – 96 h period (**Figure 4**). We found that the rate of cell proliferation is significantly reduced in PAR1-KO PC3 and significantly increased in PAR2-KO PC3 in comparison to WT PC3 cells (**Figure 4A**). On the other hand, neither PAR1 nor PAR2 antagonist had a significant effect on PCa cell proliferation (**Figure 4B**). Similarly, there was no difference seen in PAR1-KO PC3 cell proliferation rate with PAR2 agonist and antagonist treatment (**Figure 4C**). PAR1 agonist and antagonist treatment also did not show significant effects in PAR2-KO PC3 (**Figure 4D**) cell proliferation. These observations contrast with the PAR1 and PAR2 mediated response in cell migration assays and suggests that PAR1 and PAR2 have opposite roles in regulating cell migration and proliferation in PCa.

**Figure 4:**
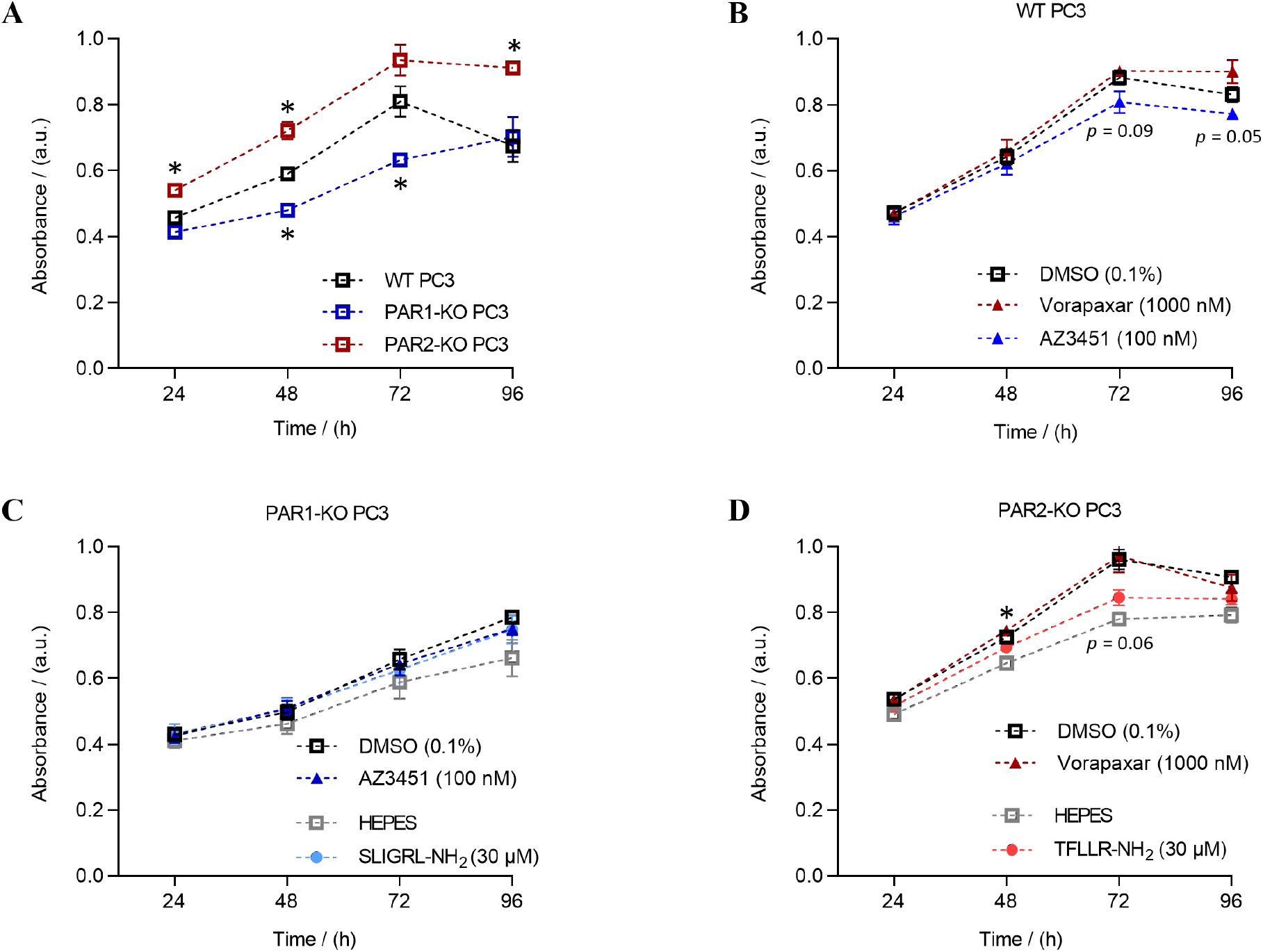
PARs regulate PC3 cell proliferation. PC3 cell proliferation assessed through a colorimetric based MTT assay over a 24-96 h period in (**A**) untreated WT, PAR1-KO and PAR2-KO PC3, (**B**) PAR1 and PAR2 antagonists vorapaxar and AZ3451, respectively, treated WT-PC3, (**C**) PAR2 antagonist and agonist SLIGRL-NH2 treated PAR1-KO PC3, and (**D**) PAR1 antagonist and agonist TFLLR-NH2 treated PAR2-KO PC3 cells. Each data point represents the mean ± SEM (N ≥ 3). Bonferroni-Dunn multiple t-tests was utilized to determine statistical significance at \**p* < 0.05 relative to the (untreated) control.

### Differentially expressed genes in PAR1-KO and PAR2-KO PC3 cells

Transcriptome profiling on total RNA extracted from WT, PAR1-KO and PAR2-KO PC3 cells was done using the Affymetrix PrimeView Human Gene Expression Array. This identified 560 (126 up-regulated and 434 down-regulated) and 2515 (1545 up-regulated and 970 down-regulated) differentially expressed genes in PAR1-KO PC3 and PAR2-KO PC3, respectively, with greater than 2-fold-change compared to WT PC3 cells (**Figure 5A**). Interestingly, PAR1 (F2R) was upregulated (fold-change 5.34) in PAR2-KO PC3 cells (**Figure 5B**). This was consistent with the increase in PAR1 mediated calcium signal seen in PAR2-KO PC3 relative to WT PC3 cells (**Figure 1A**, **B**). Several proteases including MMP1, MMP13, ADAM12 and plasminogen activator urokinase (PLAU) were downregulated in PAR1-KO PC3 cells while the serine protease inhibitor SERPINA1 was upregulated in PAR1-KO PC3 (**Figure 5C(i)**). In contrast, PAR2 KO-PC3 cells showed upregulation of several MMPs (MMP1, MMP7, MMP10, MMP13 and MMP16) and protease inhibitors (SERPINB2, SERPINE1, SERPINF1, SERPINH1, SERPINI1, SPINK5 and SPINK13) (**Figure 5C(ii)**).

**Figure 5:**
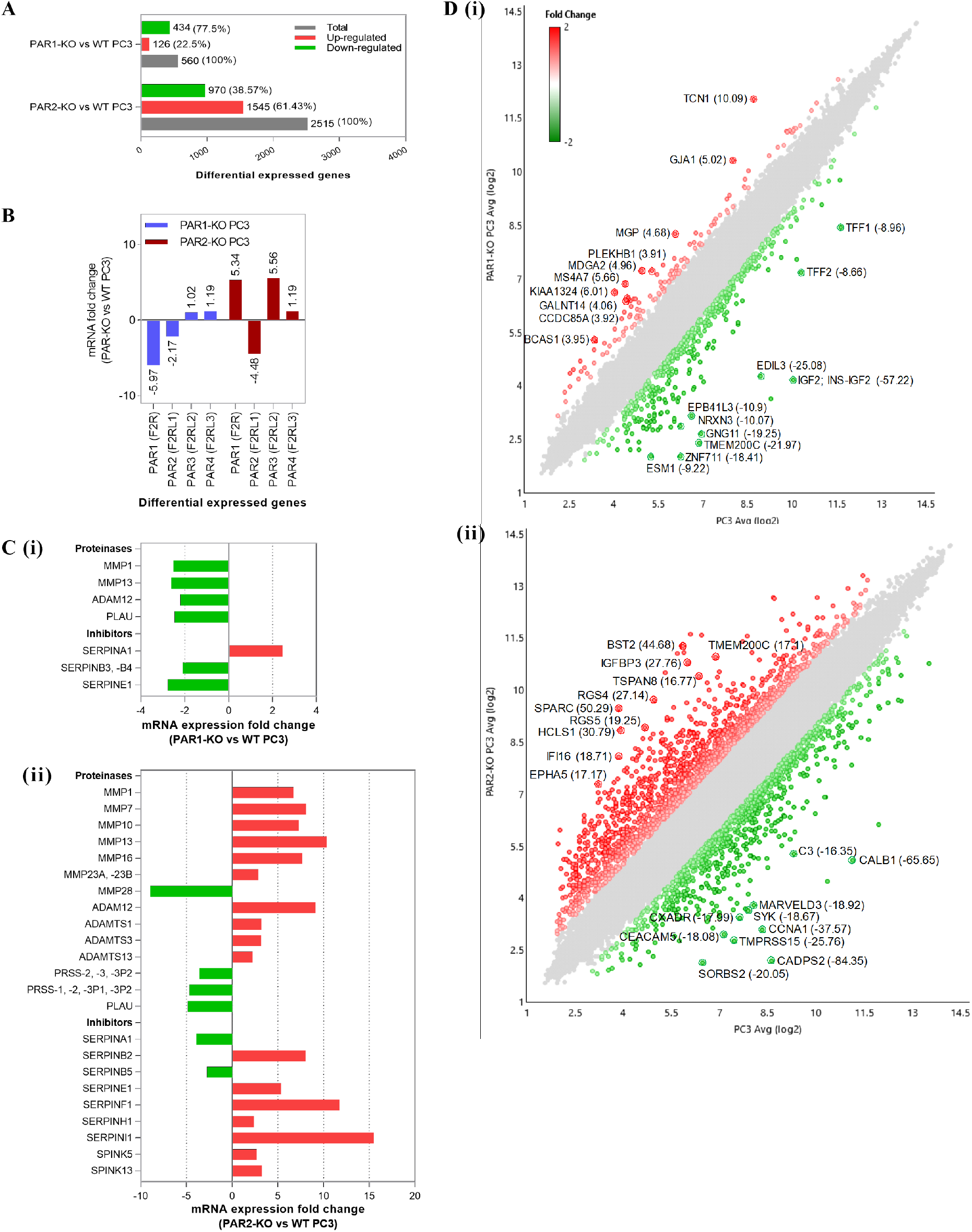
Transcriptome profiling of differentially expressed genes in PAR1- and PAR2-KO versus WT PC3 cells. Differentially expressed (**A**) total genes, (**B**) PARs, (**C**) endogenous proteinases and inhibitors, and (**D**) the top ten upregulated and downregulated genes, in PAR1-KO and PAR2-KO PC3 with greater than 2-fold-change relative to WT PC3 cells analyzed on Transcriptome Analysis Console (Affymetrix) software.

We further analyzed the top ten upregulated and downregulated genes in PAR1-KO (**Figure 5D(i)**) and PAR2-KO (**Figure 5D(ii)**) PC3 cells and examined their known roles in PCa or other cancers. As summarized in **Table S1**, tumor suppressor genes including TCN1^38^, KIAA1324^39^, MDGA2^40^ and MGP^41, 42^ were upregulated in PAR1-KO PC3 cells while GNG11^43^, ZNF711^44^ and EPB41L3^45^ were downregulated. MS4A7^46, 47^, GJA1 (Connexin 43)^48^, GALNT14^49^, BCAS1^50, 51^, and PLEKHB1^52^ genes that promote metastatic properties such as cell differentiation, migration and invasion were also overexpressed, while several PCa promoting genes including IGF2^53^, EDIL3^54^, NRXN3^55^, ESM1^56^, TFF1^57, 58^ and TFF2^57, 58^ were significantly downregulated in PAR1-KO PC3 cells.

In contrast, tumor suppressor genes including IGFBP3^59^, RGS4^60^, RGS5^61^, IFI16^62^ and EPHA5^63^ were overexpressed, and SORBS2 (ArgBP2)^64, 65^, MARVELD3^66^ and CXADR^67^ were downregulated in PAR2-KO PC3 cells. Tumor promoter genes such as SPARC^68^, BST2^69^, HCLS1^70^ and TSPAN8^71^ genes were upregulated PAR2-KO PC3, while CADPS2^72, 73^, CALB1^74^, CCNA1^75^, TMPRSS15^76^, SYK^77^, CEACAM5^78^ and C3^79, 80^ genes that are overexpressed in metastasis were significantly downregulated. Overall, this suggests that PAR1 and PAR2 regulate both tumor promoting and suppressing roles by regulating several key oncogenes associated with PCa.

Biological pathway analysis of differentially expressed genes in PAR1-KO PC3 (**Figure 6A**) and PAR2-KO PC3 (**Figure 6B**) using the WikiPathways database and manual annotation based on previous studies^81,82^ revealed roles in major PCa signalling pathways, including aryl hydrocarbon receptor, bone metamorphosis, Wnt/β-catenin, glucocorticoid receptor, immunity cell, Gαs, hypoxia, inducible nitric oxide synthase (iNOS), transforming growth factor (TGF)-β, apoptosis, cell cycle, inflammatory response, G protein and prostaglandin synthesis regulation.

**Figure 6:**
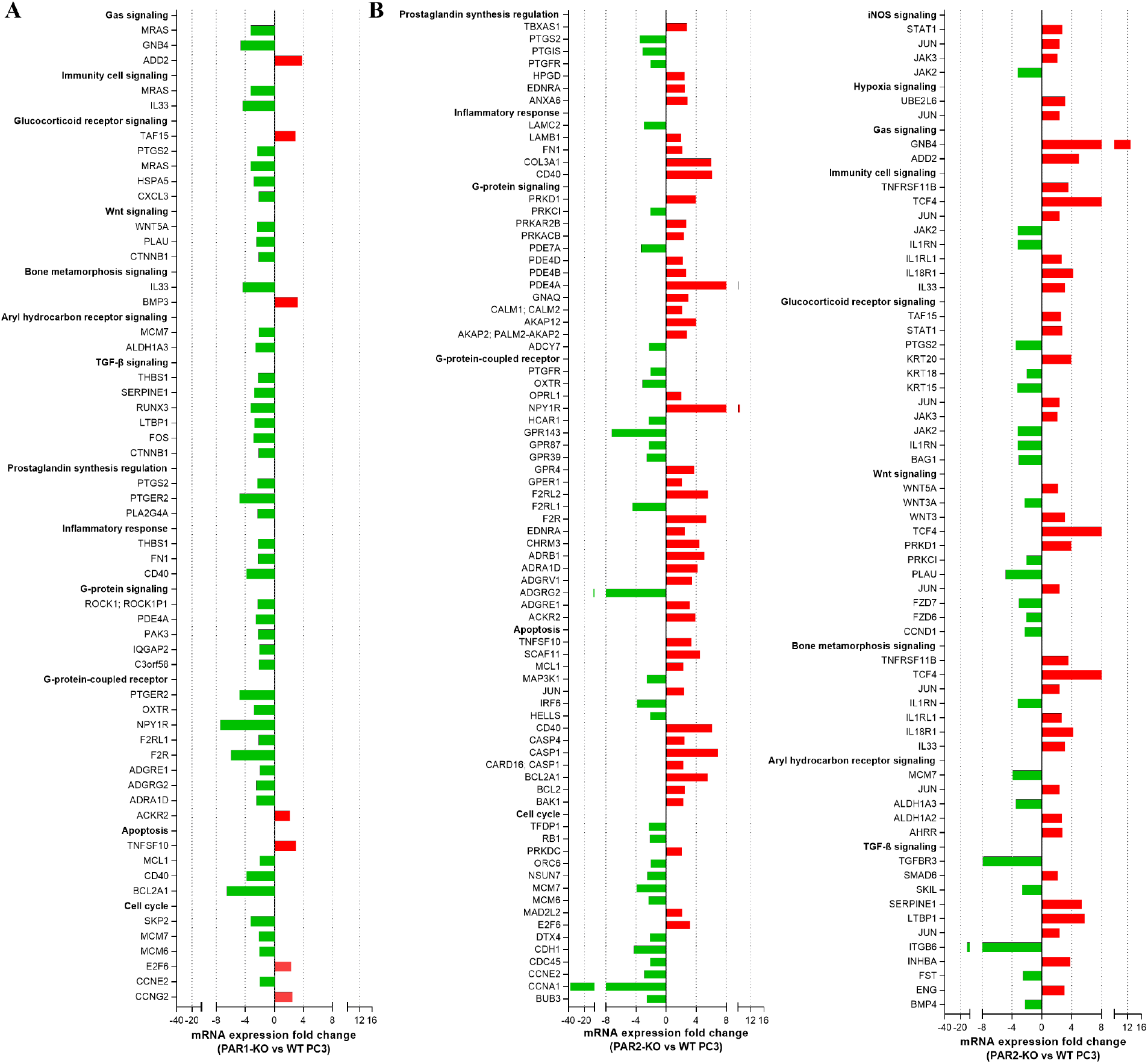
Identification of major PCa associated signaling pathways with differential expression. (**A**) PAR1-KO PC3 and (**B**) PAR2-KO PC3 with greater than 2-fold-change relative to WT PC3 cells. Pathways were analyzed and annotated by WikiPathways on Transcriptome Analysis Console (Affymetrix) software or manual annotation.

## DISCUSSION

PARs have been studied in the context of cancer in various in vitro and in vivo models, however the physiological activators of PARs remain poorly identified. In many different cancer cells, including PCa, overexpression of PAR1 and 2 is shown to participate in cancer cell motility and metastasis^83^, and postulated to have a role in tumorigenesis. As an organ that produces several proteases, including prostate-specific antigen (PSA)^84^, homologous kallikrein-related peptidase 2 (hK2)^84^, prostate/hK4^85^, and transmembrane serine protease 2 (TMPRSS2)^86^, that are known or potential activators of PARs, the prostate gland has been recognized as an organ that could undergo regulation through PAR signaling. The serine protease thrombin is also reported to enhances the expression of uPA (PLAU) through stimulation of thrombin-specific receptor in PC3 cells^87^. uPA is an initiator of pericellular proteolysis of ECM and has a role in tumor invasion, growth and metastasis. The activation of PARs in PCa by proteinases produced by the cancer cells themselves could thus serve as an important autocrine mechanism involved in cancer progression. Indeed, migration, invasion, metastasis and angiogenesis are major tumor promoting roles ascribed to proteinases in the tumor microenvironment^88^.

By harnessing the unique mechanisms of action of PAR activation involving the proteolytic cleavage of the receptor by enzymes, we were able to generate genetically encoded reporter constructs that provide information on PAR cleavage without the need for monitoring downstream effector interaction and second messenger signalling. Using these reporters, we identified elevated levels of PAR cleaving enzymes in the PC3 PCa cell line. In general PAR activation on expressing cells occurs through paracrine production of activating proteases, as for example seen in the setting of tissue injury where coagulation cascade enzymes can activate platelet expressed PARs. The autocrine activation of PARs in PC3 cells highlights yet another mechanism for activation and signaling through these receptors in disease.

Upregulation of PAR1 is closely linked to invasive phenotype and distant metastasis in several cancer cell lines^24,25^. In addition to proteinases, we identified several growth factors, cytokines and adhesion molecules as PAR1 downstream targets in PC3 cells. PAR1 primarily enhances cell invasiveness by increasing adhesion to ECM and through breakdown of ECM and basement membrane by proteinases^24^. Similarly, in several cancer cells, activation of PAR2 by its agonist peptide dramatically enhances cell migration, whereas PAR2 knockdown inhibits cell motility^29^. Furthermore, blocking of PAR2 activity with antibodies can effectively suppress tumor growth in vivo^29^.

In the case of PCa cells we found that PAR1 deficiency promoted cell migration while treatment with a PAR2 antagonist modestly but significantly inhibited cell migration. Somewhat surprisingly a PAR1 antagonist did not phenocopy the response seen in PAR1-KO PC3 cells. This could be explained in part by probe-dependence of the PAR1 antagonist, which is known to be effective in inhibiting PAR1 responses to thrombin and TFLLR-NH_2_ but may not block the MMP driven signaling through PAR1 that is implicated in PC3 cells. In contrast we see that the PAR2 negative allosteric modulator AZ3451 modestly decreased PC3 cell migration, an effect that was not observed in the PAR2-KO PC3 cells. It will be interesting to examine whether compensatory changes that allowed cells to maintain their migratory capacity where present in the PAR2-KO cells but were not engaged following acute treatment with AZ3451.

In our microarray data, key cancer associated molecular signalling pathway genes were differentially expressed when PAR1 or PAR2 was knocked out in the PC3 cell line. Several of these deregulated genes are recognized to promote PCa development and progression in humans. Consistent with previous studies performed in different cancer cell lines^20–23^, our results here support the notion that PAR1 and PAR2 signaling is directly or indirectly linked to PCa development. However, it is still unclear if PAR1 or 2 activation or inhibition is protective or detrimental in prostate cancer development since known tumor enhancing genes and tumor suppressor gene levels were affected in the PAR-KO PC3 cells. In addition, we see differential regulation of migration and proliferation when PAR activity is perturbed in PC3 cells. These differences seen in our functional studies is however not entirely unprecedented. For example connexin 43 (GJA1) overexpression increases metastatic potential in prostate cancer cell lines through promoting migration but not cell proliferation^48^. Indeed, regulation of connexin-43 gene, GJA1, may explain the enhanced migration in the absence of effects on cell proliferation seen in PAR1-KO PC3 cells, where GJA1 is upregulated.

Further assessment of PAR1 and PAR2 regulated intracellular signalling pathways in PCa is critical to fully understanding underlying mechanism of disease development and progression and in identifying specific strategies for future therapeutic intervention.

## Abbreviations

PCa: prostate cancer
PAR: Proteinase-Activated Receptor
MMP: matrix metalloproteinases
ECM: extracellular matrix
KO: knock out

## Conflicts of Interest

The authors declare no conflict of interest.

## Authors Contributions

R.R. and A.C. designed the study, performed the experiments, contributed to data interpretation, and wrote the manuscript.

## Acknowledgements

This study was supported by a Prostate Cancer Canada and Canadian Institutes of Health Research (CIHR) grants to R.R.

## SUPPLEMENTARY INFORMATION

### Supplementary figures

**Figure S1:**
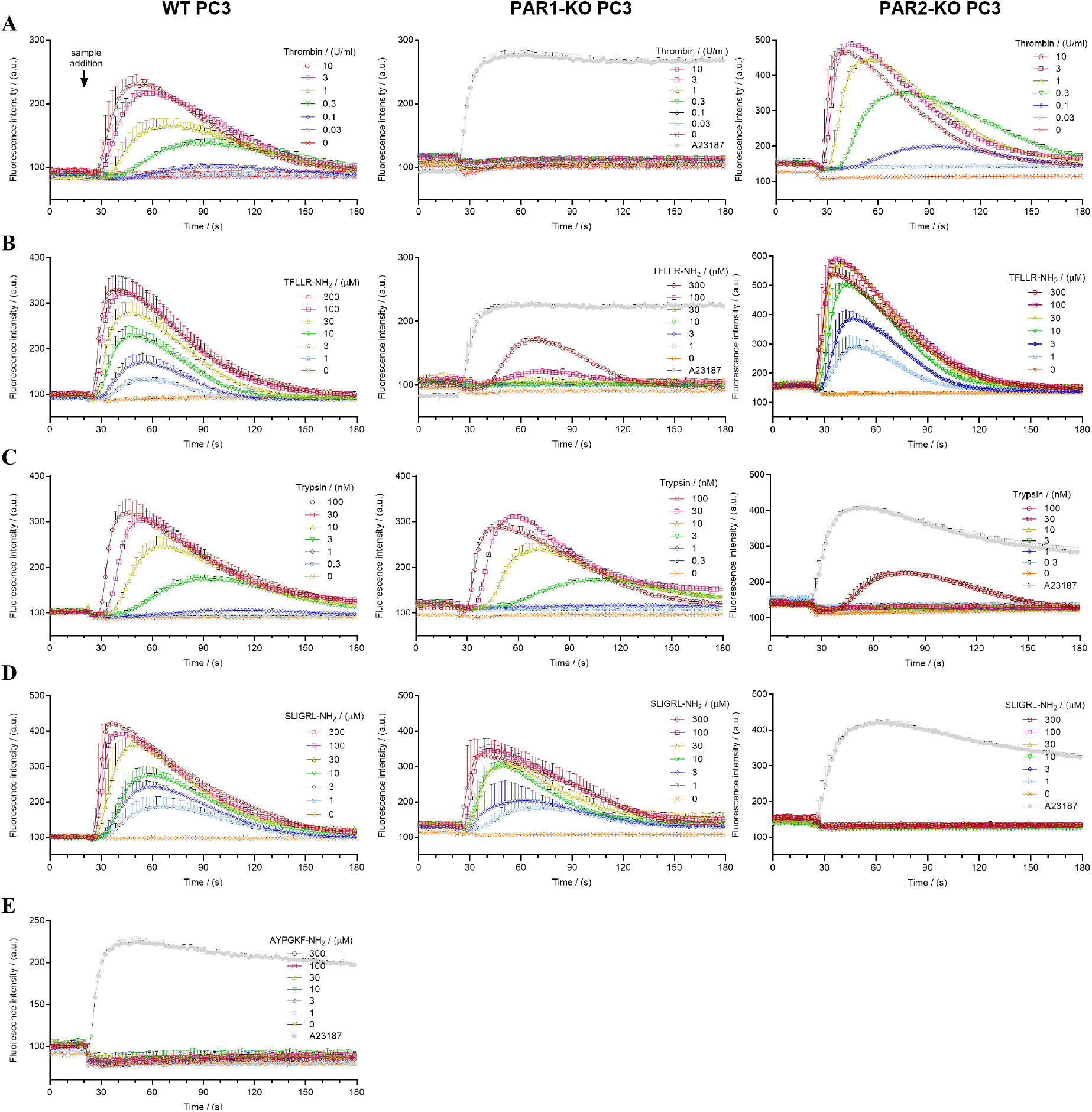
Representative traces of calcium signalling in WT, PAR1-KO, and PAR2-KO PC3 cells treated with PAR1 agonists (**A**) thrombin (0.03 – 10 U ml^-1^) and (**B**) TFLLR-NH2 (1 – 300 µM), PAR2 agonists (**C**) trypsin (0.3 – 100 nM) and (**D**) SLIGRL-NH2 (1 – 300 µM), and PAR4 agonist (**E**) AYPGKF-NH2 (1 – 300 µM).

**Figure S2:**
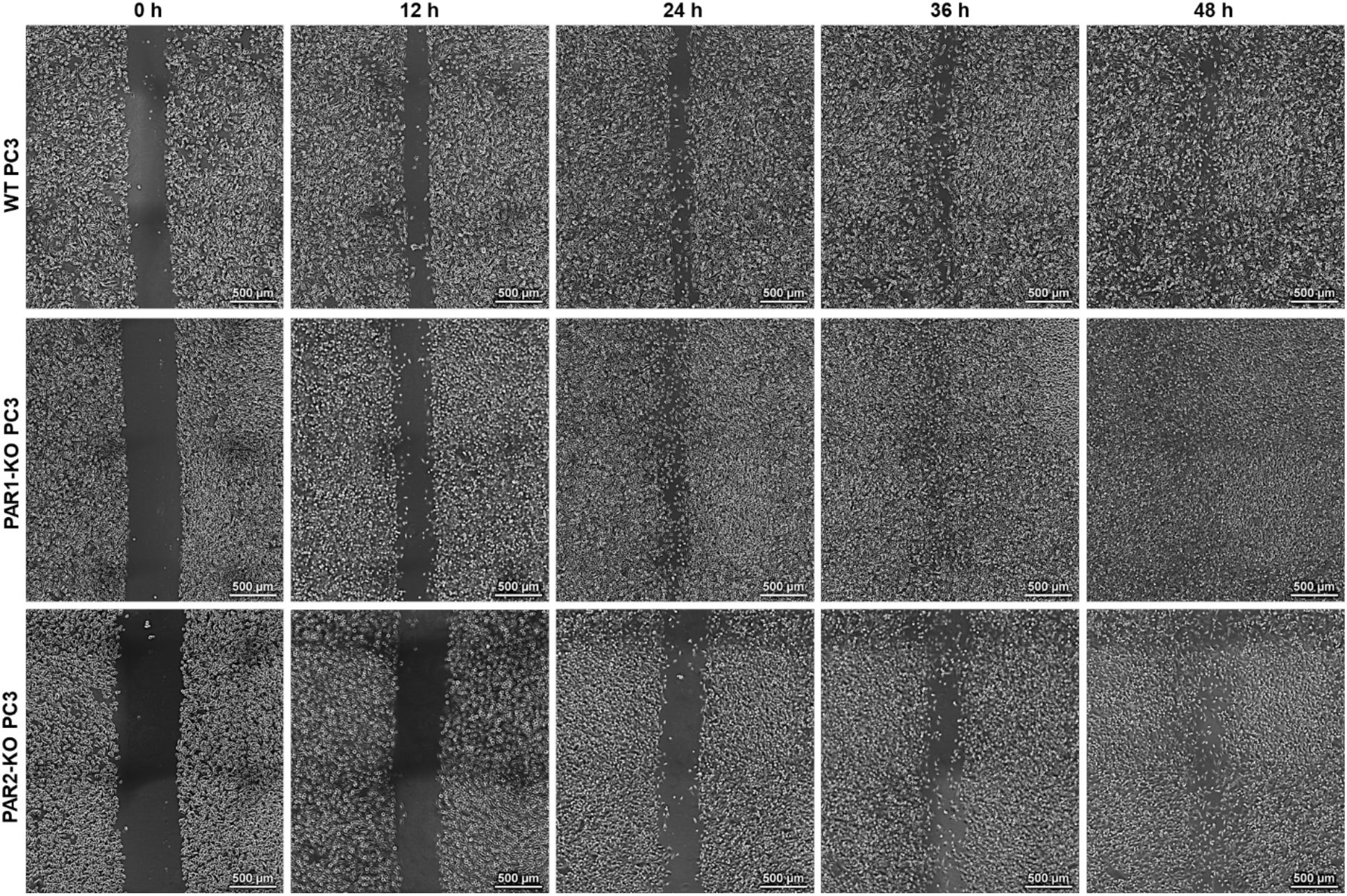
Representative images of cell migration in WT, PAR1-KO, and PAR2-KO PC3 cells over a 48 h period.

**Figure S3:**
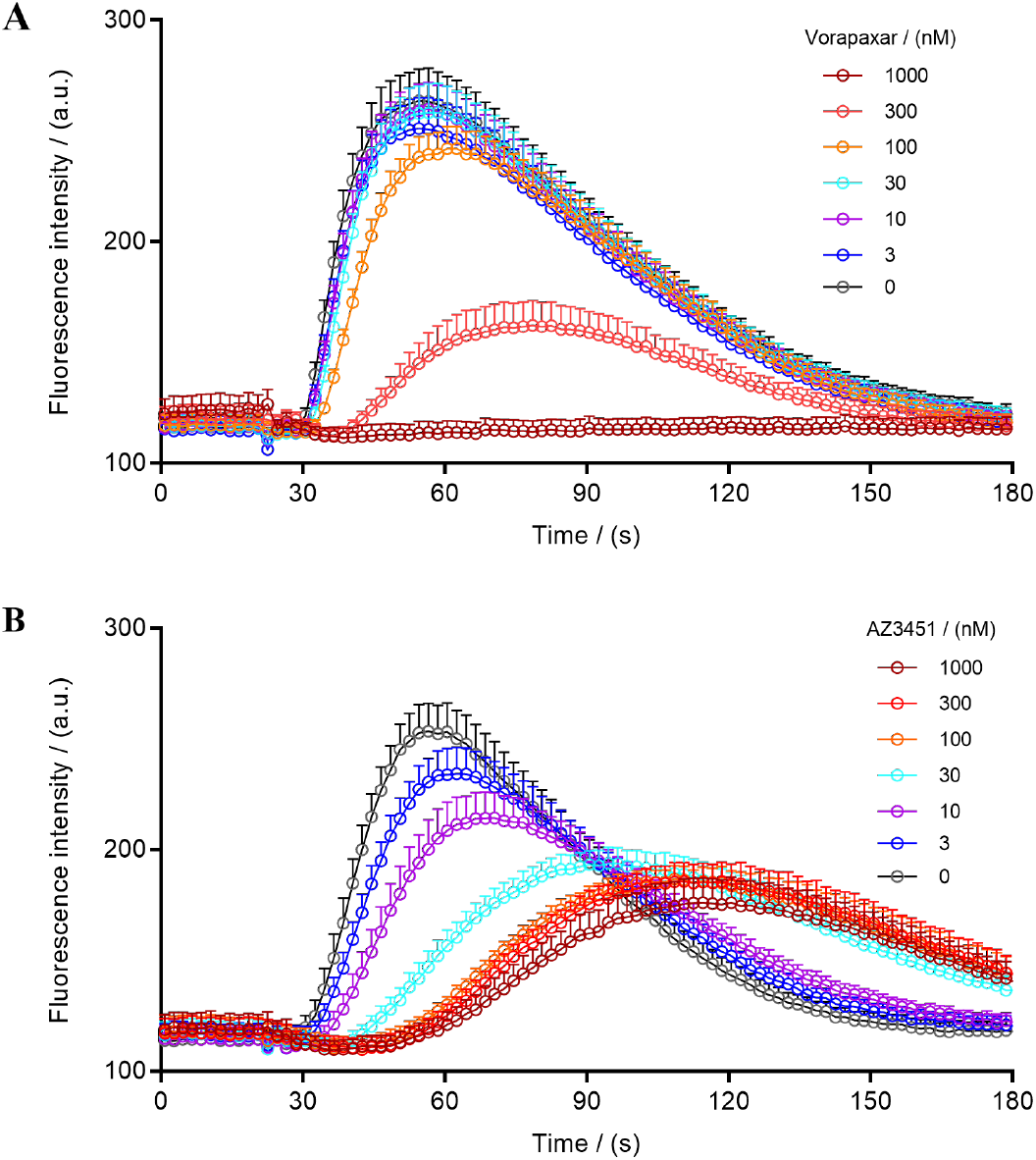
Representative traces showing inhibition of PAR1 and PAR2 calcium signaling responses to thrombin (1 U ml^-1^) and trypsin (10 nM) respectively in the presence of PAR1 and PAR2 selective antagonists vorapaxar and AZ3451 in PC3 cells.

**Table S1:**
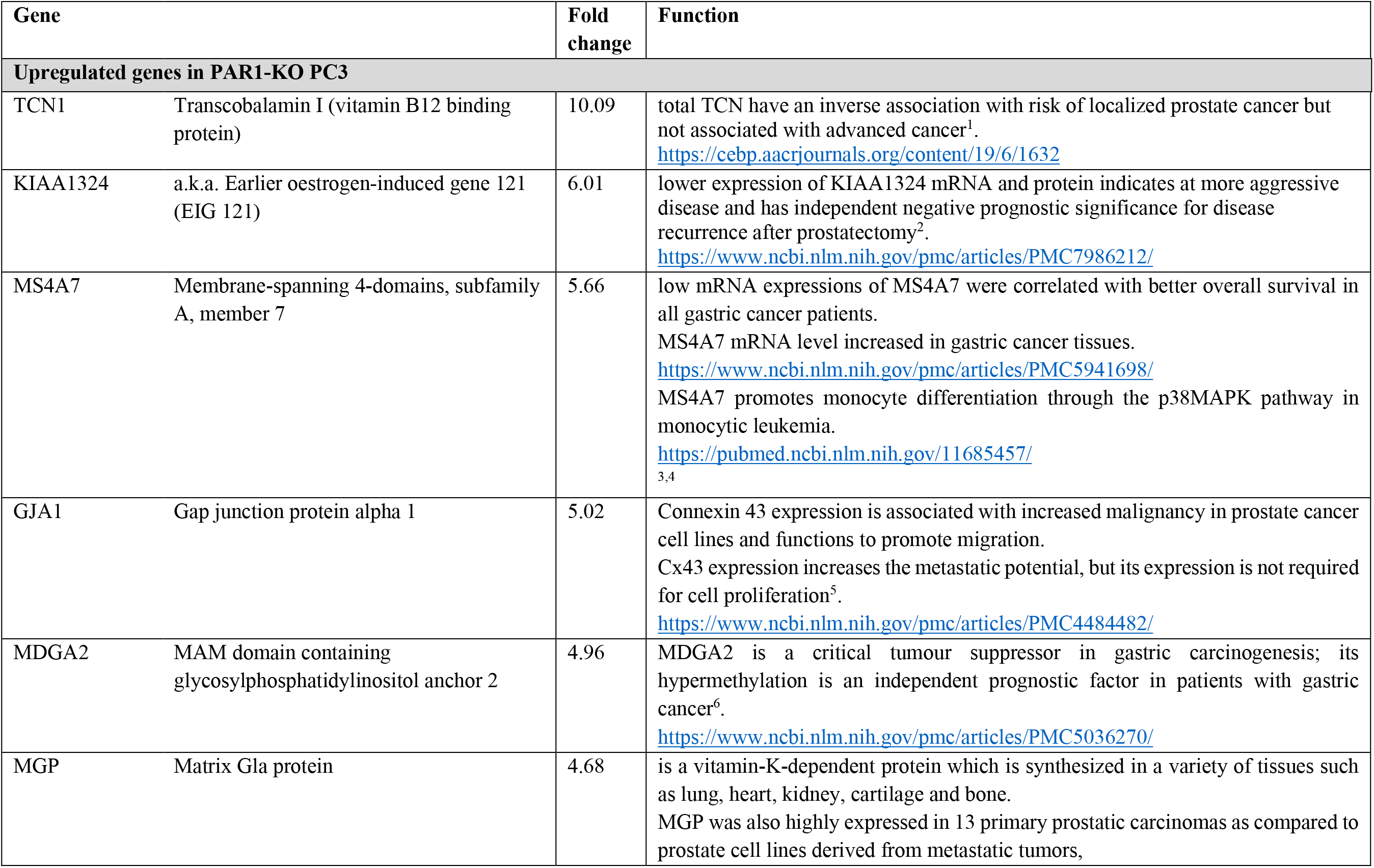
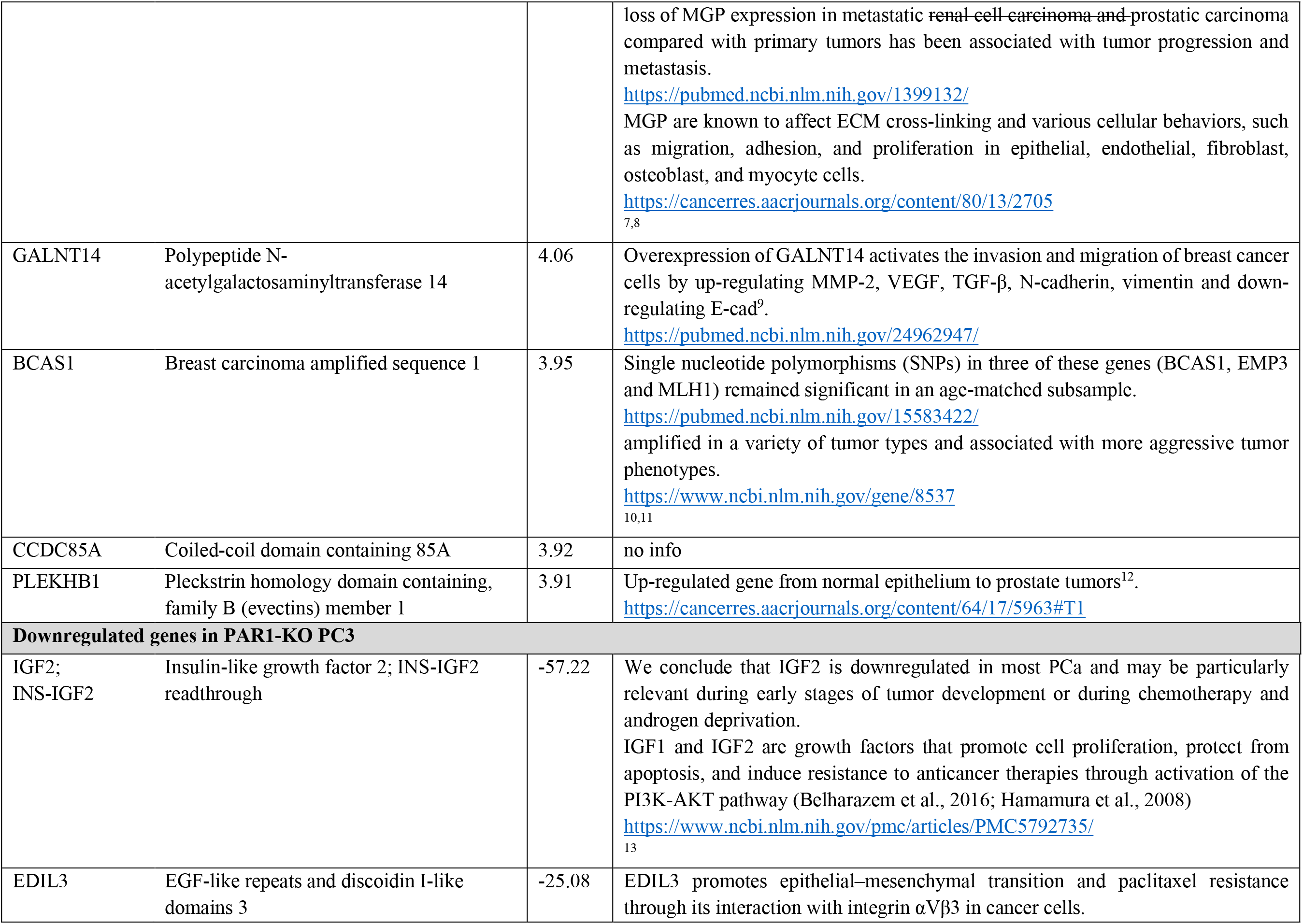
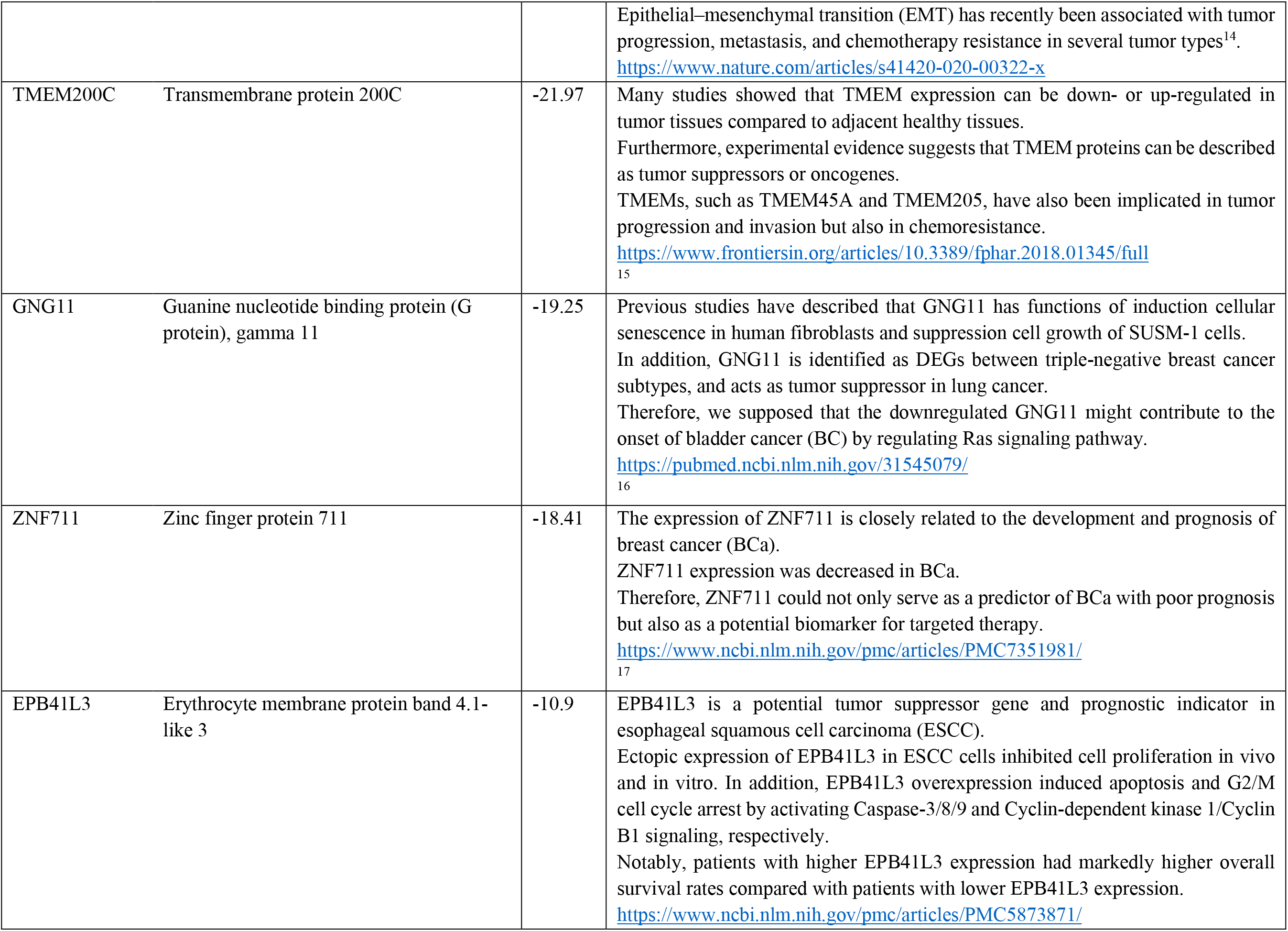
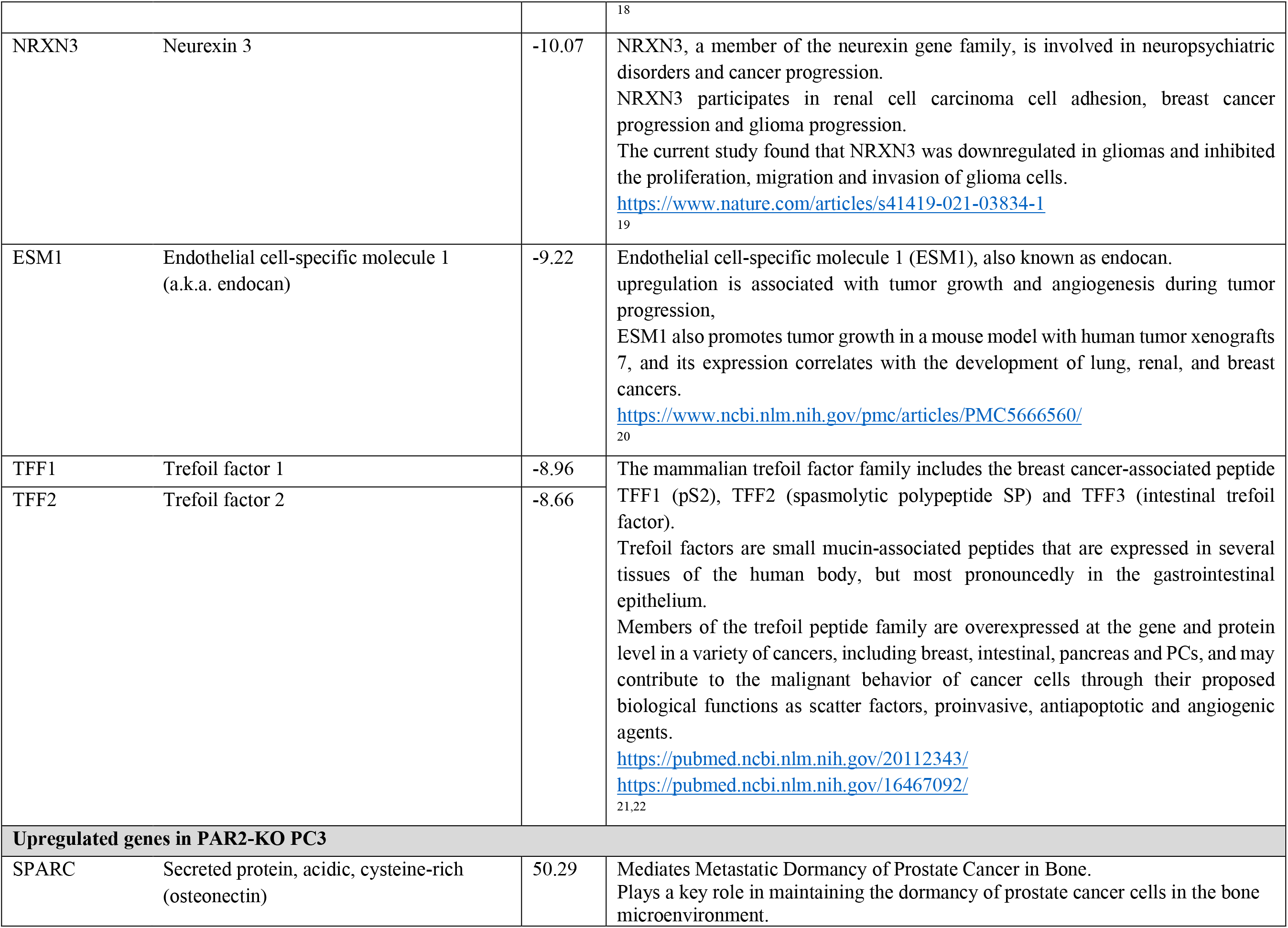
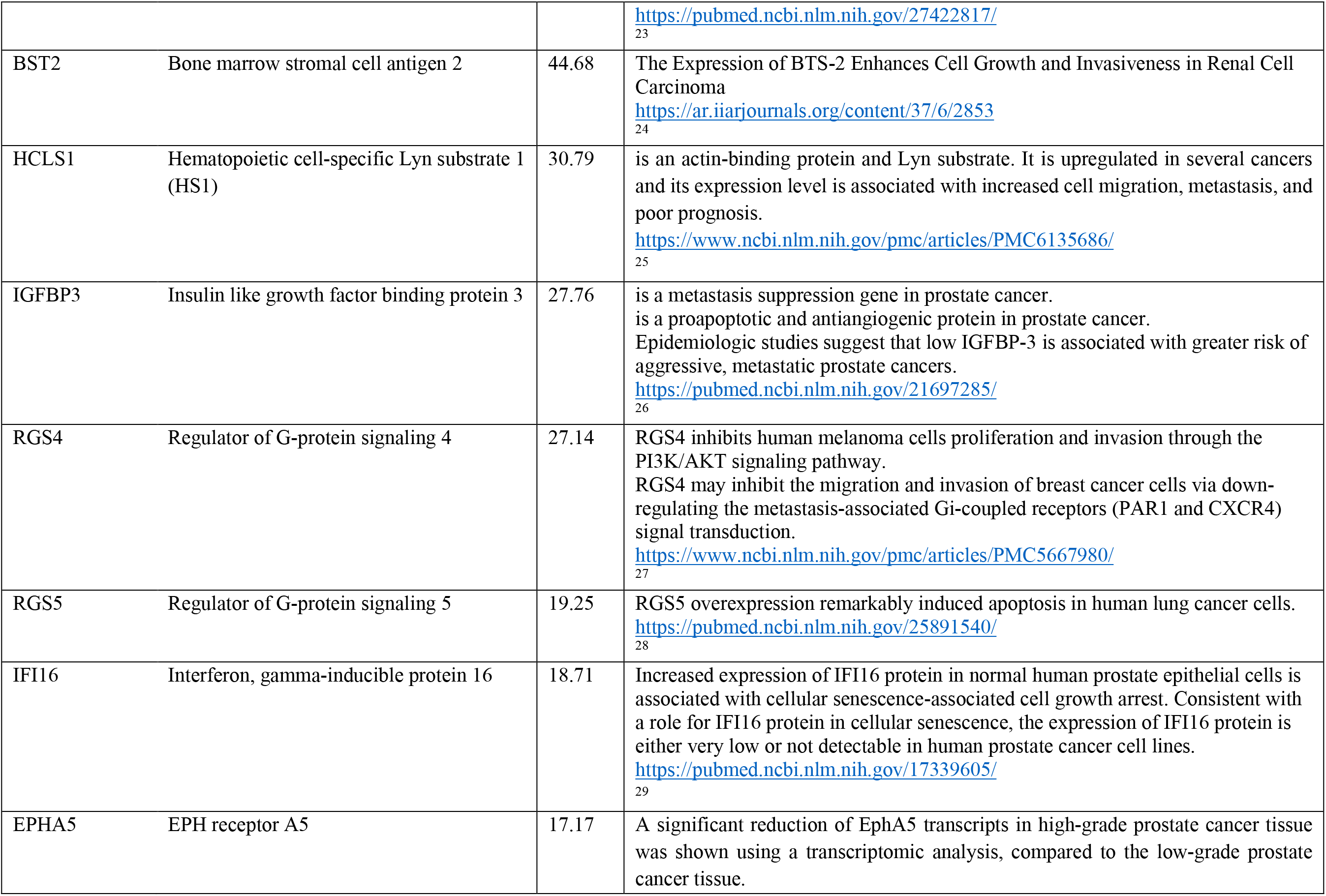
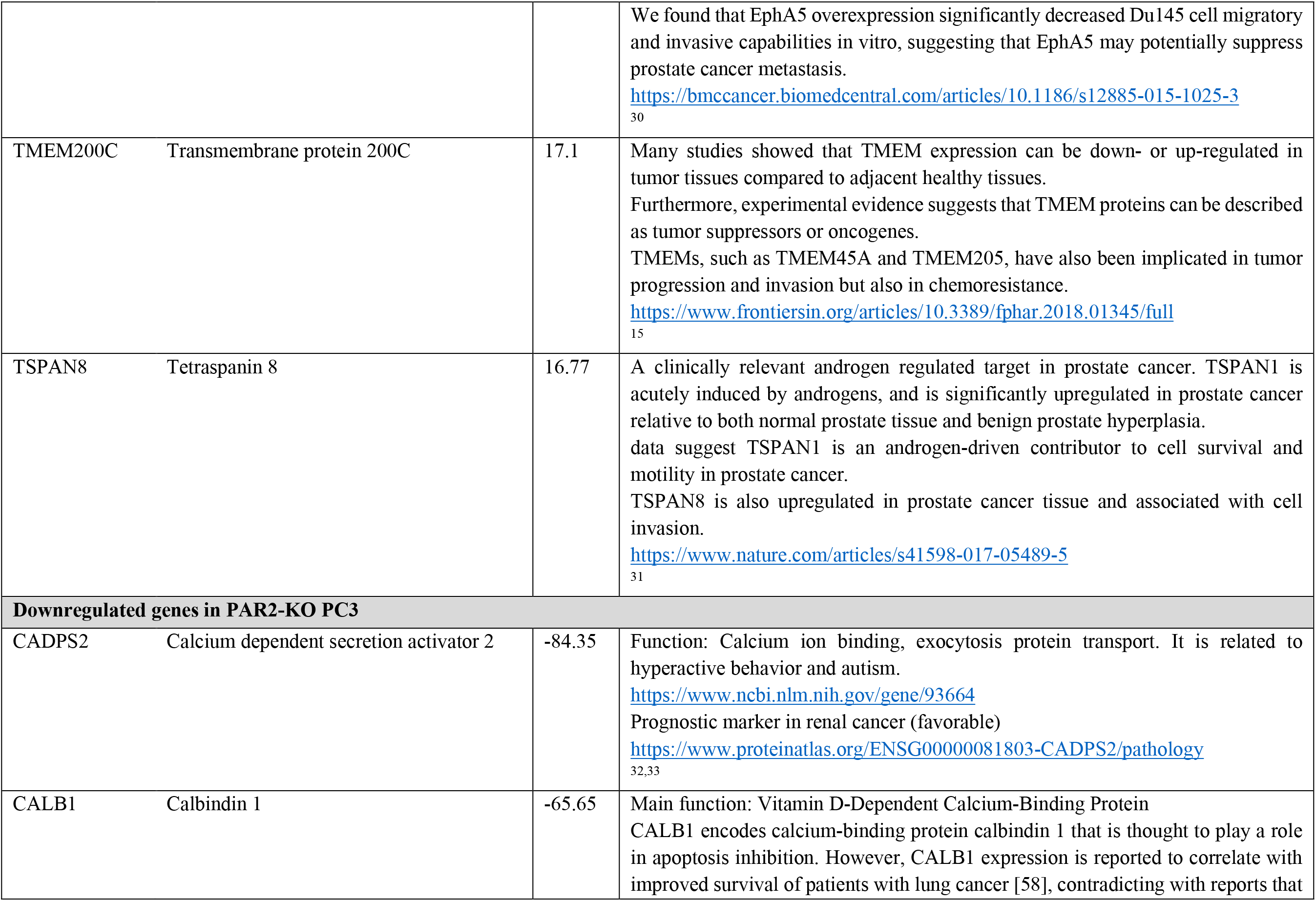
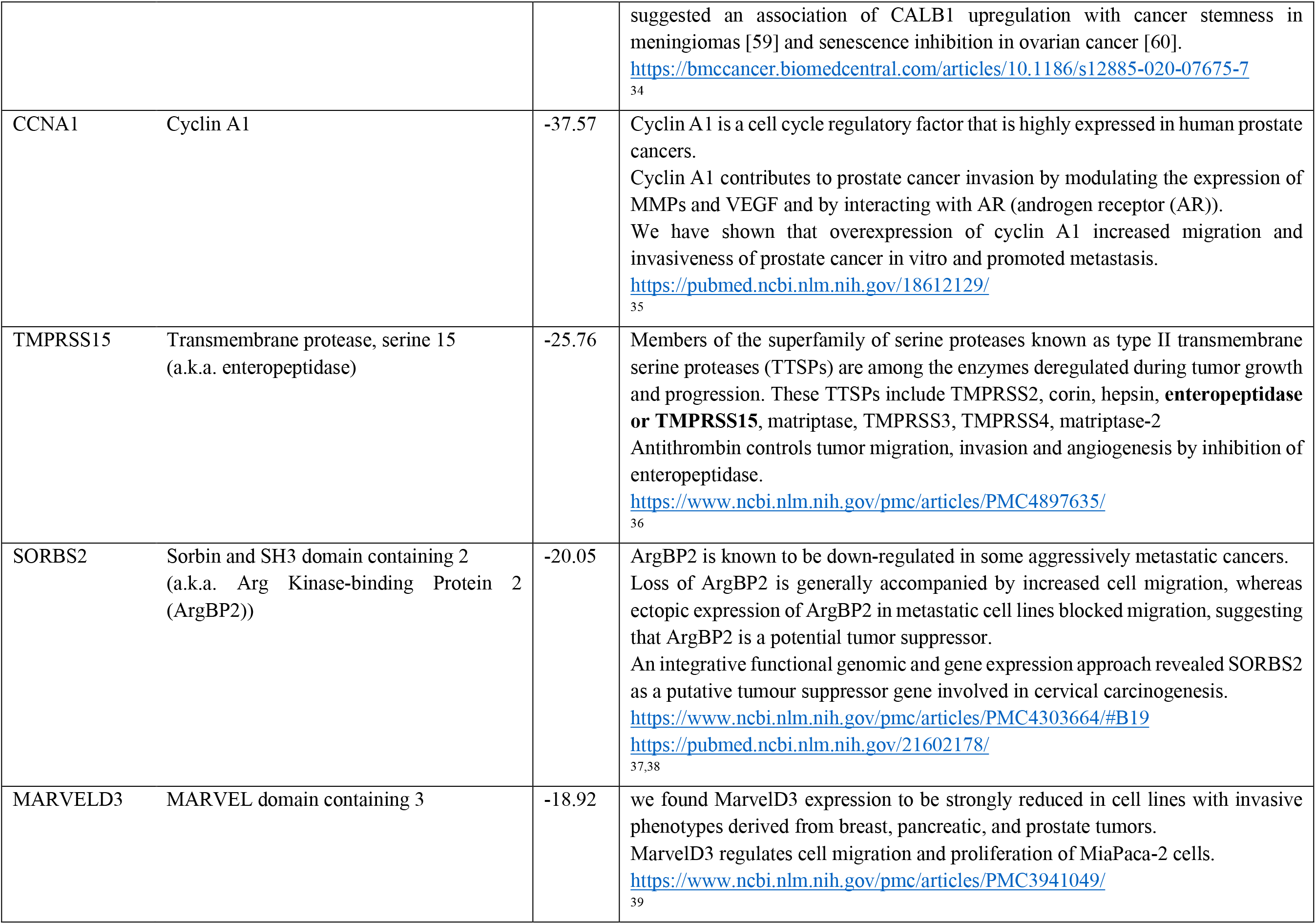
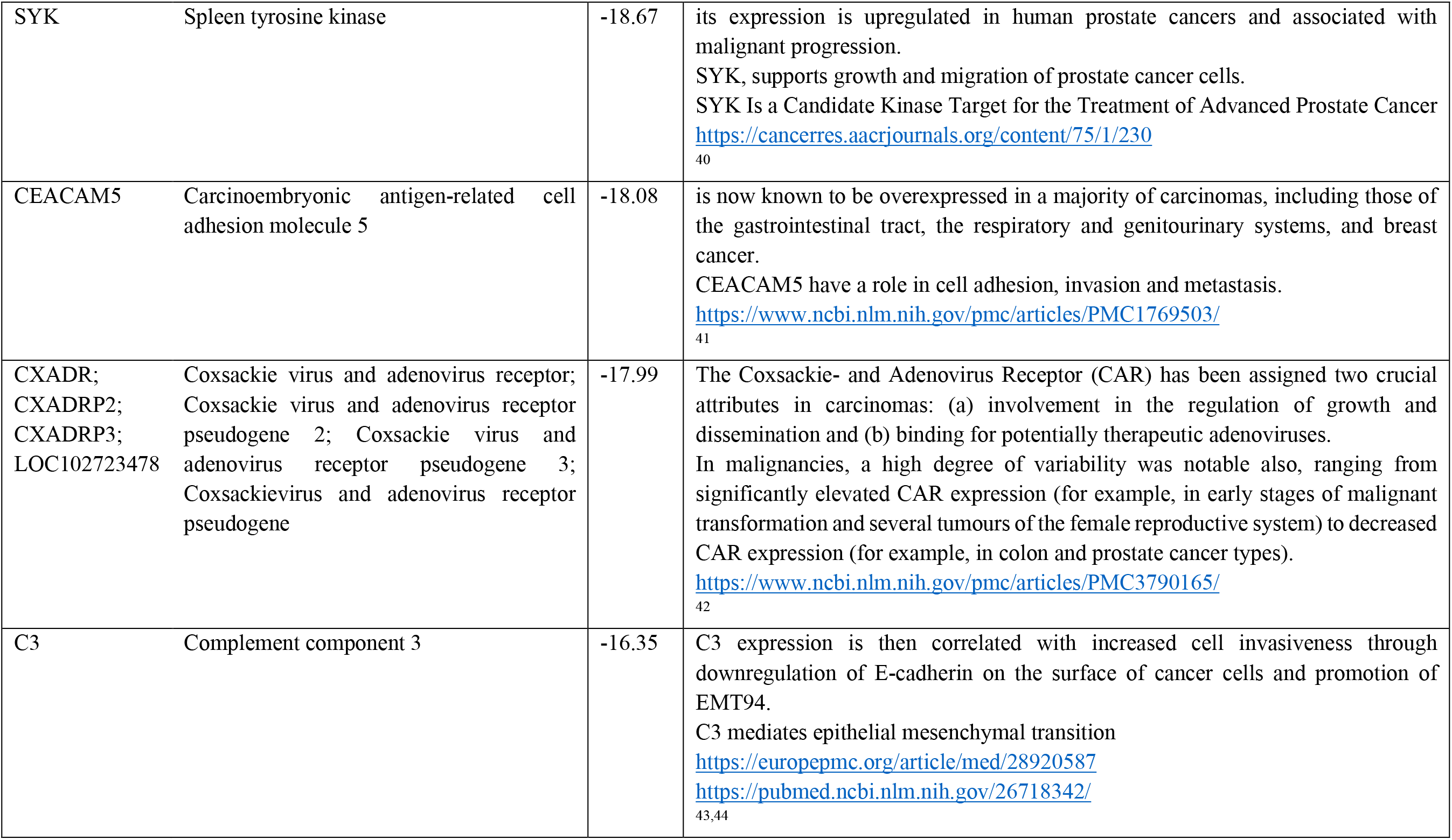
Top ten upregulated and downregulated genes in PAR1-KO PC3 and PAR2-KO PC3 versus WT PC3 cells.

## Notes

### Competing Interest Statement

The authors have declared no competing interest.

